# Local Mean Suppression Filter for Effective Background Identification in Fluorescence Images

**DOI:** 10.1101/2024.09.25.614955

**Authors:** Bogdan Kochetov, Shikhar Uttam

## Abstract

We present an easy-to-use, nonlinear filter for effective background identification in fluorescence microscopy images with dense and low-contrast foreground. The pixel-wise filtering is based on comparison of the pixel intensity with the mean intensity of pixels in its local neighborhood. The pixel is given a background or foreground label depending on whether its intensity is less than or greater than the mean respectively. Multiple labels are generated for the same pixel by computing mean expression values by varying neighborhood size. These labels are accumulated to decide the final pixel label. We demonstrate that the performance of our filter favorably compares with state-of-the-art image processing, machine learning, and deep learning methods. We present three use cases that demonstrate its effectiveness, and also show how it can be used in multiplexed fluorescence imaging contexts and as a denoising step in image segmentation. A fast implementation of the filter is available in Python 3 on GitHub.

## 1 Introduction

Fluorescence microscopy images of biological tissues exhibit high complexity, which is characterized by diverse factors such as spatial heterogeneity of tissue architecture, crowded cells, complex morphology, noise, and varying contrast between background and the foreground components [1]. As a result, despite considerable work in this area, precise identification of the image background in fluorescence microscopy remains a non-trivial problem in many biological contexts [1]. Indeed, the list of methods for segmenting images into foreground and background proposed over the last half century is long [1, 2, 3, 4, 5, 6, 7] and includes a variety of conceptual approaches that include histogram-based and adaptive thresholding, active contours, region merging, Markov models, clustering, graph cuts, watershed transform, total variation, and wavelet analysis. Beyond these primarily unsupervised image processing and machine learning approaches, deep learning methods have achieved significant success in image segmentation over the past decade using deep learning (DL) models [8, 9, 10]. In particular, many of these unsupervised [11, 12, 13, 14, 15, 16, 17, 18, 19, 20, 21] and DL [22, 23, 24, 25, 26, 27, 28, 29, 30, 31, 32, 33] methods have been applied to instance segmentation of cells and nuclei in microscopy images, an important step in computational and systems biology studies. Critically, the accuracy of each of these segmentation methods can benefit from a reliable pre-processing step that improves image quality by removing background noise associated with sensor noise, light scattering and diffusion, and autofluorescence. Therefore, new approaches have recently been proposed to suppress the background in fluorescence microscopy images [34, 35]. Moreover, a variety of background removal pipelines can be built using simple methods – for example, median and bilateral filtering, rolling ball algorithm, histogram equalization – and open access software packages [36]. However, each of these methods is usually suited to fluorescence images with specific foreground and background characteristics.

In this paper, we introduce a new method, built-upon a new local mean suppression filter (LMSF), to solve the problem of background identification and removal in fluorescence images. LMSF itself compares the intensity of a pixel with the mean intensity of its local neighborhood to label the pixel as a foreground or background pixel. Our method utilizes LMSF to generate multiple labels for the pixel as a function of the size of its local neighborhood. The labels are then aggregated by our method to generate the final label for the pixel. Varying the neighborhood size provides a natural way to account for short and long range variations in the fluorophore intensity, and decide whether the pixel is truly foreground or background. As a result, our LMSF based method is able to separate the background from the foreground in images with complex spatial morphology and architecture. Importantly, it is able to precisely identify dim foreground objects from the background with ambiguous transition regions. To the best of our knowledge no unsupervised approaches have demonstrated this ability applicable to diverse contexts. Specially trained and fine tuned DL methods do demonstrate this ability, however, it is primarily in the context of cell and nucleus segmentation, and not in general background identification scenarios. In contrast our LMSF based method is not limited to cells and nuclei but can be used to identify tissue components arising from fluorescence of fluorophores conjugated with any molecule of interest. We demonstrate this ability through multiple use cases and potential applications of the LMSF method.

The paper is organized as follows. In Section 2 we introduce LMSF for analog and digital signals, and then generalize the latter for processing fluorescence images. We next introduce our LMSF based method as well as compare its efficacy with other unsupervised and DL methods in Section 3. We then present three important use cases for the LMSF method in Section 4. In Section 5 we present application of our method to scenarios beyond background identification. Specifically, we demonstrate its use for clustering and preprocessing of multiplexed fluorescence images of tissue sections. We conclude in Section 6.

## 2 Local Mean Suppression Filter

LMSF is designed to identify background in fluorescence images [37, 38] with sharp and gradual variations in intensity due to low or high contrast of the tissue components and their dense or sparse localization in an image with acquisition noise. To achieve this goal LMSF itself performs a simple operation: compute the ratio of the pixel intensity to the mean intensity of its local neighborhood of a given size. If the ratio is below a threshold, label the pixel as a background pixel, otherwise, label it as a foreground pixel. LMSF gives the background label a value of zero, while it preserves the original intensity value for the foreground label.

We begin the formulation of LMSF for an analog signal representing the idea, properties, and parameters of LMSF.

### 2.1 Analog signal

For *s*(*t*), a real-valued function of continuous time, LMSF response *ŝ*(*t*) is given by,

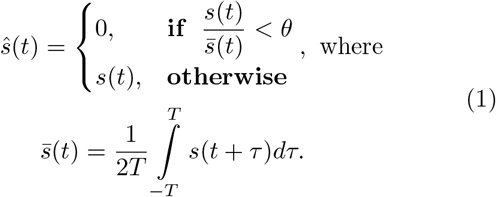

Here 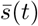 is the local mean computed within the time window defined by the parameter *T*, and *θ* is the threshold parameter. Eq. (1) states that at time instant *t*, LMSF replaces the original signal value *s*(*t*) with 0 only if the ratio of the instantaneous signal value *s*(*t*) to its local mean 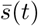 is less than the threshold parameter *θ*.

The threshold parameter *θ* should not be set too low because lower the threshold value, higher the probability of falsely labeling a low signal level (background) as a high signal level (foreground). On the other hand, setting the parameter *θ* to be high creates the opposite fallacy of falsely labeling a high signal level as a low signal level. To motivate selection of *θ* value, we consider a sine-squared signal, *s*(*t*) = sin^2^(*t*), that mimics a signal with varying intensity. Its local mean value is,

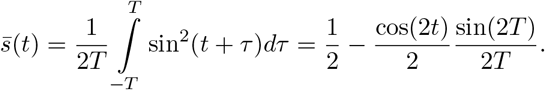

As can be seen, this local mean value fluctuates around a value of 0.5. Therefore, setting *θ* = 0.5, will result in LMSF labeling values of *s*(*t*) approximately less than 0.25 as the low signal level (Fig. 1). As we show in this paper, *θ* = 0.5 turns out to be a good default value for real-world examples with the optimal range being 0.5 ⩽ *θ* ⩽ 1.

**Figure 1.**
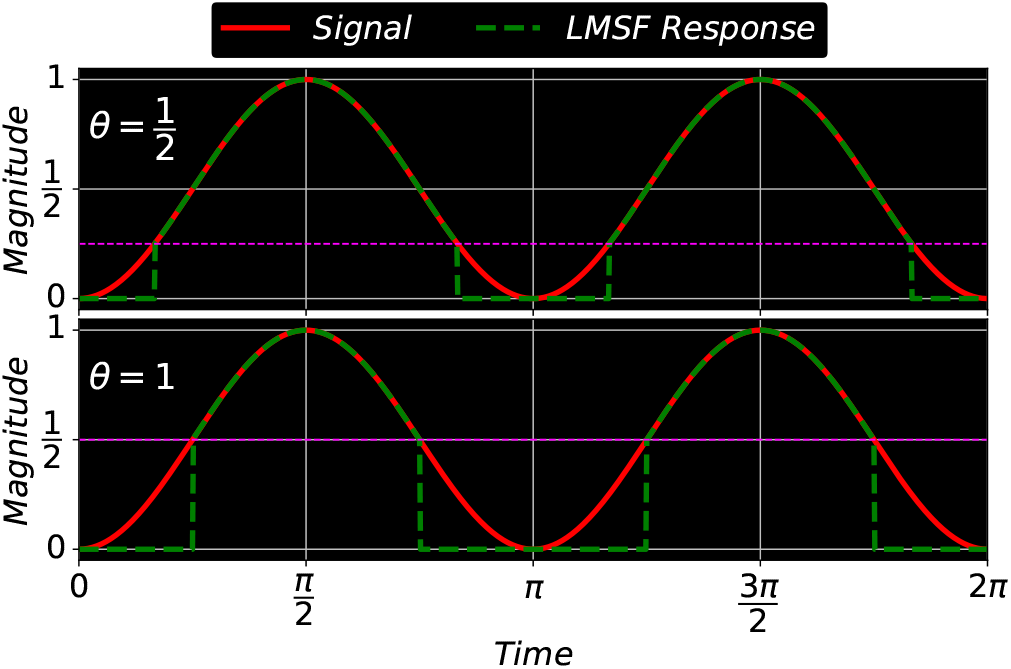
A sine-squared signal and two LMSF responses (1) for *θ* = 0.5 (upper panel) and *θ* = 1 (lower panel). In both cases 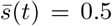 because the length of averaging window is exactly equal to an integer number of the signal periods, i.e. *T* = *kπ/*2, *k* ∈ N. Setting *θ* to 0.5 (or 1.0) results in LMSF replacing the signal values below 0.25 (or 0.5) with zeros, as indicated by the green dashed line in the upper (lower) panel.

### 2.2 Digital Signal

Sampling the analog signal *s*(*t*) – usually implemented via Nyquist sampling – results in a sequence of equidistant signal samples {*s*_*k*_}, where *s*_*k*_ = *s*(*k*Δ*t*), Δ*t* is the period of sampling, *k* = 1, …, *K*, and *K* is the number of samples. The corresponding discrete LMSF response is given by,

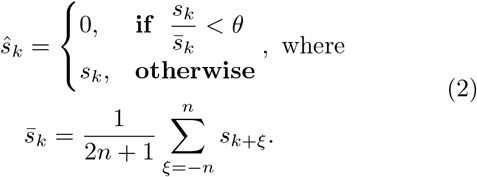

We note that in Eq. (2) the length of averaging window, defining the neighborhood around *k*-th sample, is specified by the parameter *n*.

A numerical example illustrating the LMSF response (2) as a function of *n* is illustrated in Fig. 2. It plots a digital signal (red solid line) consisting of 71 samples and the LMSF responses (green dashed line) corresponding to *n* = 2 (upper panel), *n* = 5 (middle panel), and *n* = 20 (lower panel). In all three cases *θ* = 0.5. This example illustrates how LMSF response changes as a function of *n*, and how it can be used to identify different patterns of transition from a low to a high signal level. For example, using LMSF with narrow window size (*n* = 2 and 5), we identify narrow gaps between high level signal samples. On the other hand, LMSF with a wider window (*n* = 20) allows us to identify wider gaps as well as regions of gradual signal transition from a high to a low level. However, an excessive increase in *n* results in two filtering effects which can be used to perform an effective background identification. First, narrow gaps are no longer identified. Second, the filtering of wide low level (background) regions saturates at certain value of the parameter *n* (lower panel in Fig. 2).

**Figure 2.**
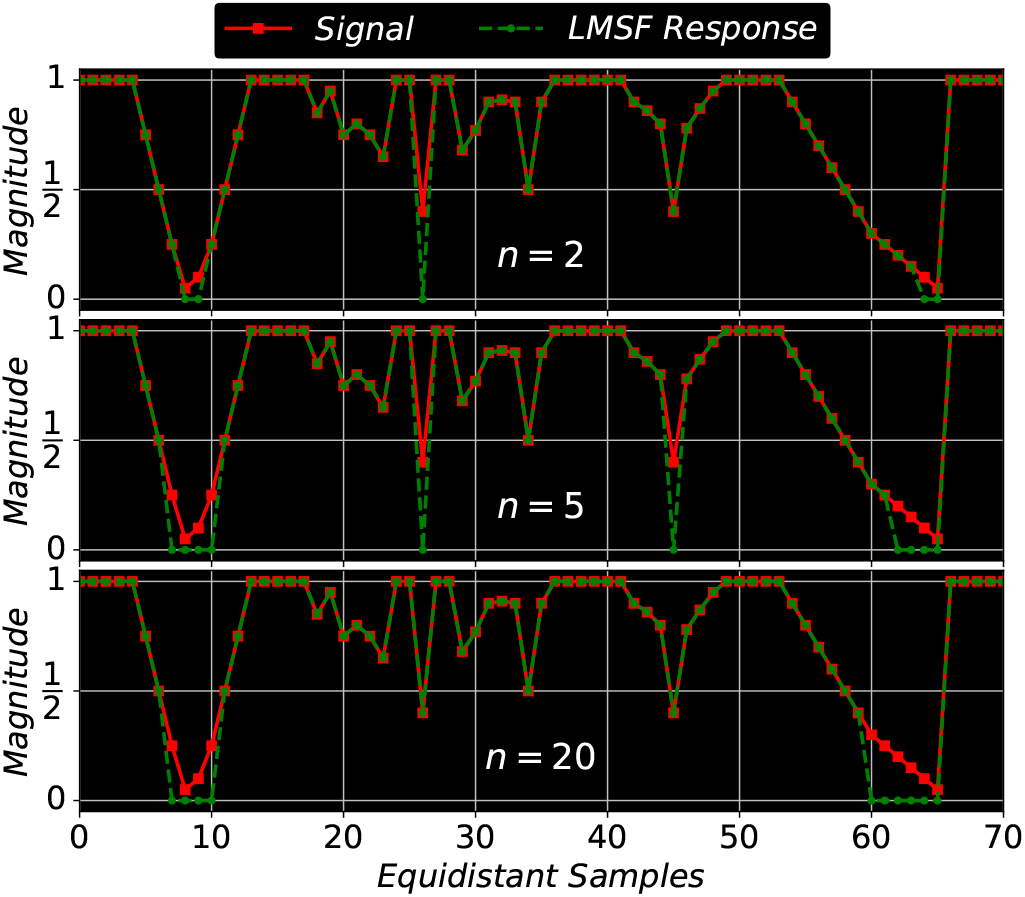
A digital signal and three LMSF responses (2) for *n* = 2 (upper panel), *n* = 5 (middle panel), and *n* = 20 (lower panel). In all cases *θ* = 0.5.

As shown in Fig. 2, the LMSF response (2) always has the same length (number of samples) as an input signal sequence. We achieve this by averaging only the available signal values at neighborhoods of the end points of a finite signal sequence. For example, to calculate the local mean at the left (right) end point we sum *n* + 1 signal values taken from the right-hand (left-hand) neighborhood of the end point and then divide the resulting sum by *n* + 1. Similarly, to calculate the local mean at the second left or right point we sum *n* + 2 available signal values taken form the corresponding neighborhood of the point and then divide the resulting sum by *n* + 2. Likewise, we apply such averaging with a shorter window to all neighborhoods of the end points of the finite sequence, where the available signal values do not overlap completely with the full size (2*n* + 1) averaging window. This adaptive averaging does not include any padding of unavailable signal values that allows us to avoid the occurrence of numerical edge effects due to signal padding (Fig. 2). The adaptive averaging is equally applicable for small and large averaging window lengths.

### 2.3 Digital Image

LMSF (2) for digital signals can straightforwardly be extended to two dimensions, applicable to images acquired on monochrome image sensors. Such an image can be considered as a double sequence of equidistant signal samples, represented as *I*(*k, m*), containing *K* rows (*k* = 1, …, *K*) and *M* columns (*m* = 1, …, *M*) Therefore, in the two-dimensional case LMSF is defined as,

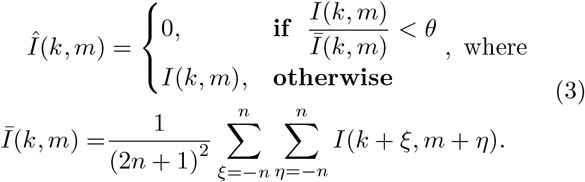

Eq. (3) states that for a given image pixel (*k, m*), LMSF replaces the original intensity value, *I*(*k, m*) with 0 only if the ratio of the pixel intensity to the mean intensity, averaged locally over (2*n* + 1) × (2*n* + 1) neighborhood of pixel (*k, m*), is below the threshold parameter *θ*. Otherwise the filter response, *Î*(*k, m*) returns the original intensity. The value of *n* is bounded between 1 and *n*_max_, where *n*_max_ is given by

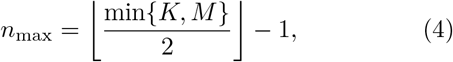

and ⌊·⌋ denotes the floor function.

## 3 Background Identification in Fluorescence Images

For many practical cases, we set *θ* = 0.5 in Eq. (3). That means, each pixel with an image intensity less than half of the corresponding local mean intensity is nullified, i.e. the original pixel intensity value is replaced by zero. Moreover, as we demonstrate in this section, by having fixed *θ* and varying the side length of the square averaging window (parameter *n*), we can effectively identify both narrow and wide gaps between complex foreground objects (clusters of cell nuclei).

### 3.1 LMSF Method for Image Background Identification

Biological tissue is heterogeneous with varying density and localization of its different components. Fig. 3a depicts an exemplar that has been imaged using Hoechst marker based fluorescence imaging to capture cell nuclei. Here spatial localization of gray intensity values – due to fluorescence expression of Hoechst marker – captures the spatial structure of cell nuclei. The 200 × 200 pixel image contains both bright nuclei with high intensity values and nuclei with very low intensity values, which make them almost indistinguishable from the background. We use it as a visual example to demonstrate the effectiveness of LMSF (3) in identifying image background. To capture the variation in nuclei intensity values and the diversity of their spatial localization we sequentially apply LMSF with increasing values of *n*, i.e. to capture the observed heterogeneity at short and long range we set *n* = 5, 10, 20, 40, 80, and *n*_max_ = 99. In all cases we set *θ* = 0.5. Figs. 3b-g illustrate these LMSF responses calculated using Eq. (3), where the identified background is shaded in blue. As can be seen, smaller values of *n* precisely capture the narrow background regions around the densely packed nuclei and their edges where the intensity changes are sharper (Figs. 3b-d), while larger values of *n* capture wider gaps and the empty regions, where intensity variations are gradual (Figs. 3e-g). The cumulative response of this sequential application – obtained via sequential logical (Boolean) multiplication of the corresponding backgrounds – is shown in Fig. 3h. The pseudocode for this cumulative LMSF response is presented in Algorithm 1. As can be seen, the LMSF method is able to precisely identify both exact edges of all foreground objects and the whole background (Fig. 3h).

**Figure 3.**
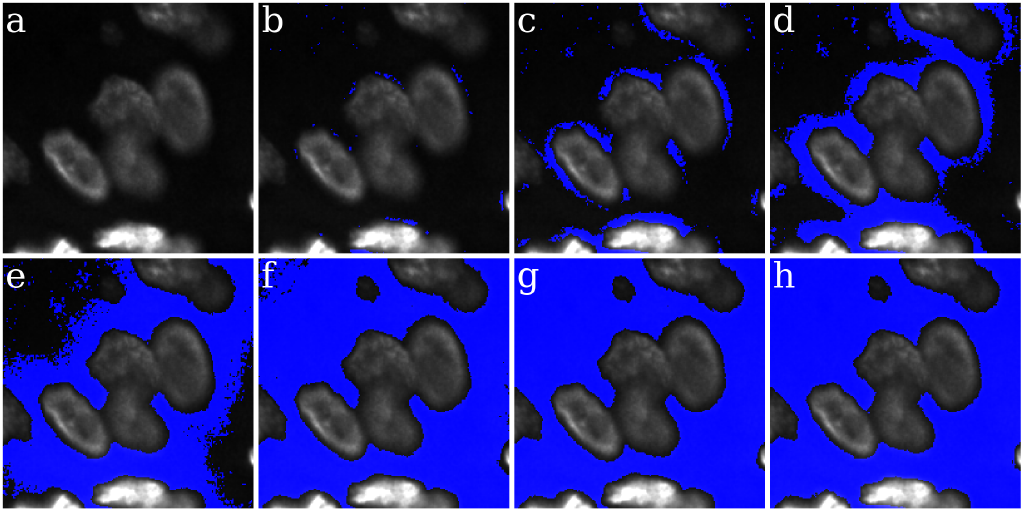
Illustration of the LMSF background identification in an image containing both bright and dim foreground objects. (a) Input image: 200 × 200 fluorescence image, where grayscale intensity indicates expression of nucleus marker (Hoechst). (b)-(g) Responses of LMSF (3) to the input image respectively corresponding *n* = 5, 10, 20, 40, 80, and 99. (h) Cumulative LMSF response (Algorithm 1) with *N* = {5, 10, 20, 40, 80, 99}. Blue color indicates the pixels having zero LMSF responses. In all cases *θ* = 0.5.

#### Algorithm 1

LMSF method for background identification in a grayscale image.

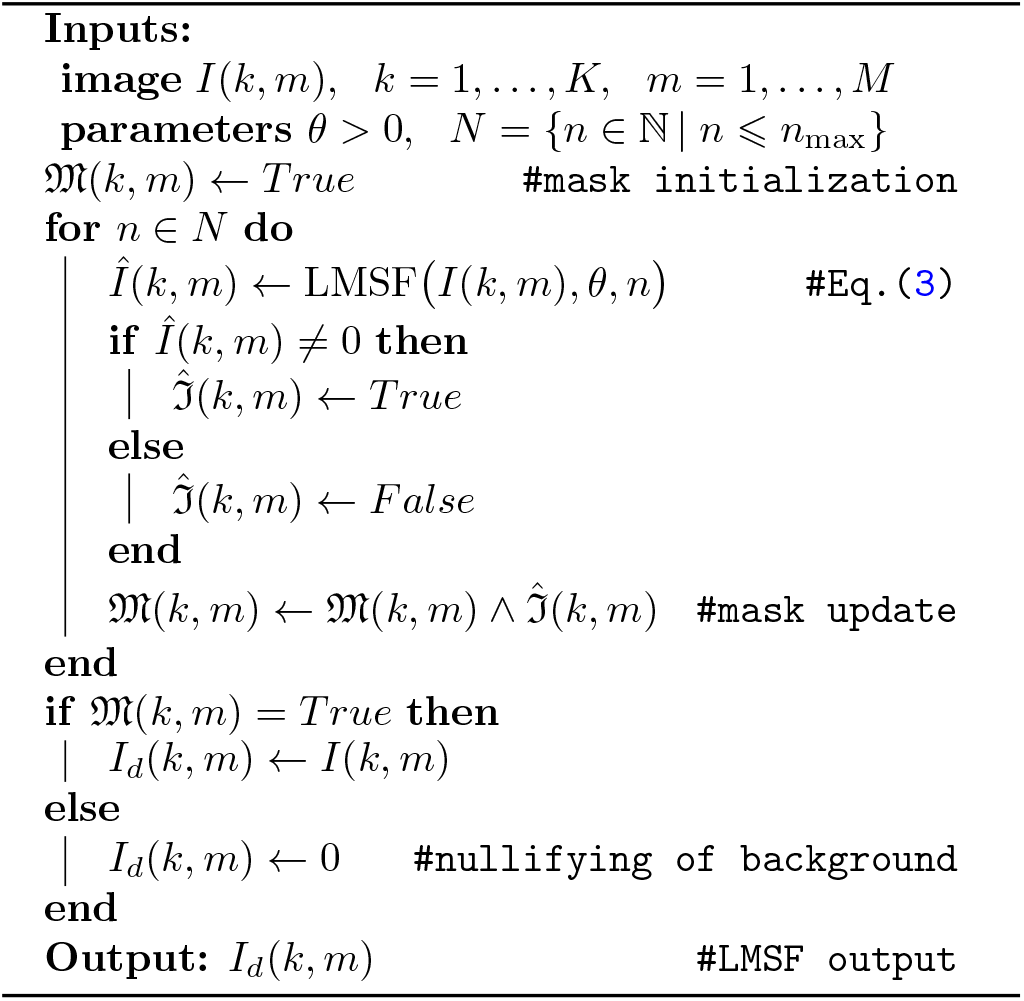

### 3.2 Comparison of LMSF with unsupervised and deep learning methods

We next compare the performance of our LMSF method with multi-level Otsu thresholding [39], Chan-Vese algorithm based on level sets [40], Li thresholding based on optimally minimizing the cross-entropy [41], and Sauvola local thresholding based on adaptive thresholding of every image pixel [42], four representative, well-known and robust unsupervised methods for binary image segmentation. We also include comparisons with Cellpose [28] and Mesmer [29], the two primary state-of-the-art supervised DL methods for performing instance segmentation of cells and nuclei in fluorescence images. To perform the comparison, we apply each of these methods to the image shown in Fig. 3a. Each subfigure in Fig. 4 shows foreground objects outlined by green lines (left panel) and the resulting binary (foreground/background) segmentation mask (right panel) predicted by each of these methods. Specifically, Fig. 4a shows the image with green outlines around foreground predicted by the lower threshold of 3-level Otsu method (left panel) and the corresponding binary segmentation mask (right panel) indicating the image foreground and background in white and black colors, respectively. Corresponding results for Chan-Vese method with *µ* = 1*/*4 and *λ*_1_ = *λ*_2_ = 1 are shown in Fig. 4b. Figs. 4c and d, respectively show the results for Li thresholding, and Sauvola local thresholding with its parameters set to *w* = 199 and *k* = 0.2.

**Figure 4.**
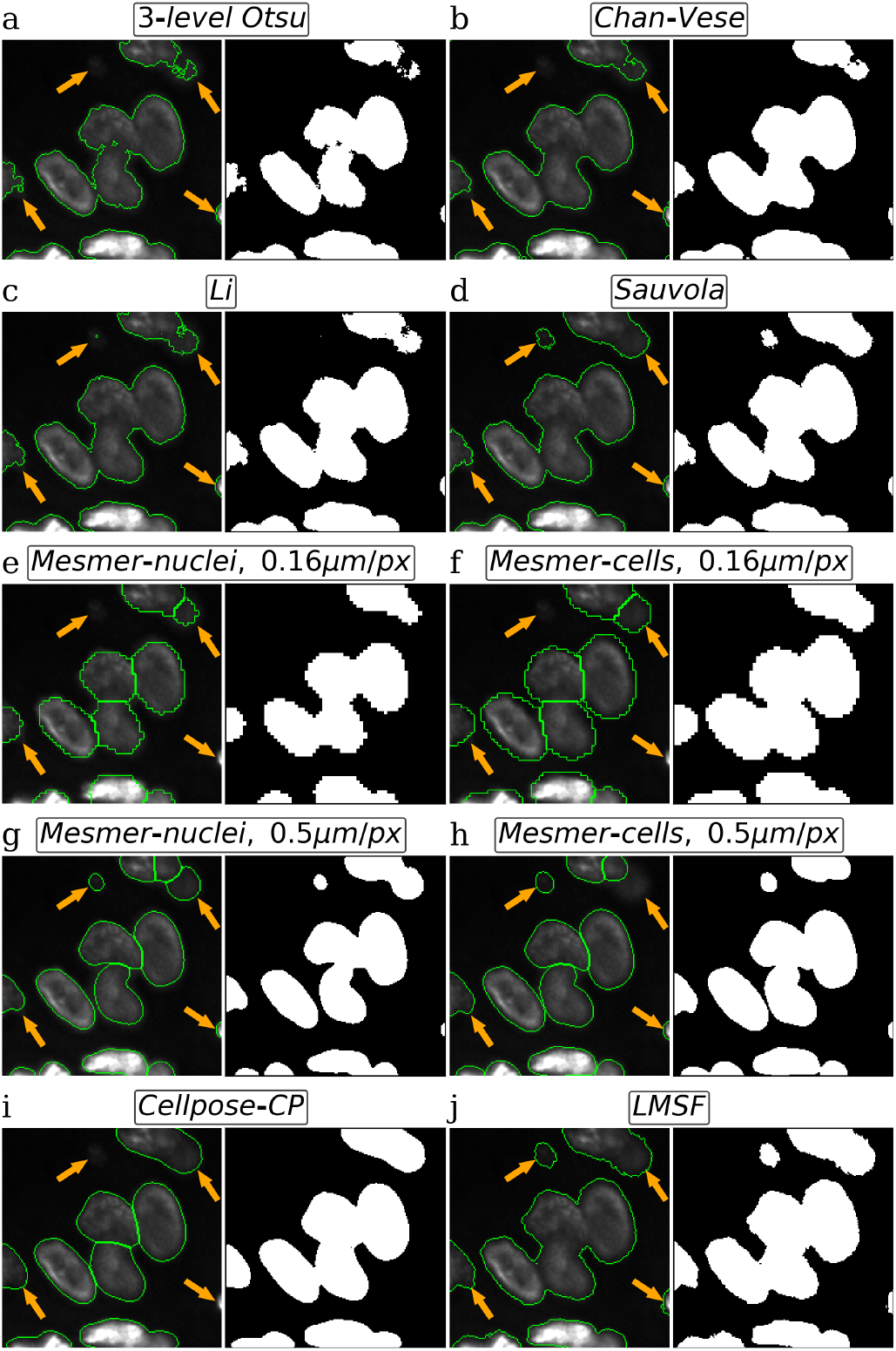
Visual comparison of image background identification for five unsupervised (Otsu, Chan-Vase, Li, Sauvola, and LMSF) and two DL (Mesmer and Cellpose) methods. For every method, using the same test image we circle green contours around foreground objects (nucleus clusters) predicted by the method and plot the corresponding binary segmentation mask, where white and black respectively indicate the foreground and background of the image. Four orange arrows point out the parts of foreground objects, which are the most difficult to identify.

When applying Mesmer method we used its Deep-Cell 0.12.6 model. We varied two Mesmer parameters: 1. `compartment`, which specifies *‘whole-cell’* (default) or *‘nuclear’* segmentation modes to predict respectively cells or nuclei; and 2. `image_mpp`, an image resolution measured in microns per pixel. Since the image (Fig. 3a) was acquired using an imaging sensor with a pixel pitch of 0.16 µm/pixel, we set this value for `image_mpp` using both *‘nuclear’* (Fig. 4e) and *‘whole-cell’* (Fig. 4f) segmentation modes. Moreover, in Figs. 4g and h we show Mesmer foreground predictions respectively for *‘nuclear’* and *‘whole-cell’* modes when image_mpp was set to its training (default) resolution, i.e. 0.5 µm/pixel. Cellpose results are shown in Fig. 4i, where we used the *CP* model from the *‘model zoo’* of Cellpose version 2.1.0. We chose *CP* model because it produced the best segmentation for the image in comparison to all other models from the *‘model zoo’*. Before predicting labels, we estimated the cell diameter using the Cellpose size calibration procedure. Finally, Fig. 4j shows the results of our LMSF method (Algorithm 1) with *θ* = 0.5 and *N* = {5, 10, 20, 40, 80, 99}. We note that Fig. 4j depicts the same result as Fig. 3h with modification to match the format of Fig. 4.

A visual comparison of the results presented in Fig. 4 suggests that global thresholding and level set methods tend to miss regions of image foreground (nuclei) with low contrast (Figs. 4a, b, and c), while specialized DL methods can miss small isolated nuclei (Figs. 4e, f, and i) as well as low contrast portions of bigger nuclei (Figs. 4e, f, h, and i). In addition, DL methods often smooth the edges of the nuclei (foreground objects) that leads to squeezing and narrowing of their actual boundaries (Figs. 4g, h, and i). On the other hand, Sauvola (Fig. 4d) and our LMSF methods (Fig. 4j), two unsupervised algorithms based on local thresholding, avoid such drawbacks. However, Sauvola method, designed for binarizing poorly illuminated or stained images [42], can effectively identify only locally homogeneous foreground objects, that is, objects having a small spread of intensity values. As a result, Sauvola method tends to exclude foreground pixels belonging to a nucleus whose intensities are relatively low, incorrectly labeling them as background. This is illustrated in Figs. 5b-d, where Sauvola method spuriously identifies large central regions of the nuclei as background (shown in blue) across varying window size parameter. On the other hand, as shown in Figs. 5e-h, our LMSF method avoids such drawbacks.

**Figure 5.**
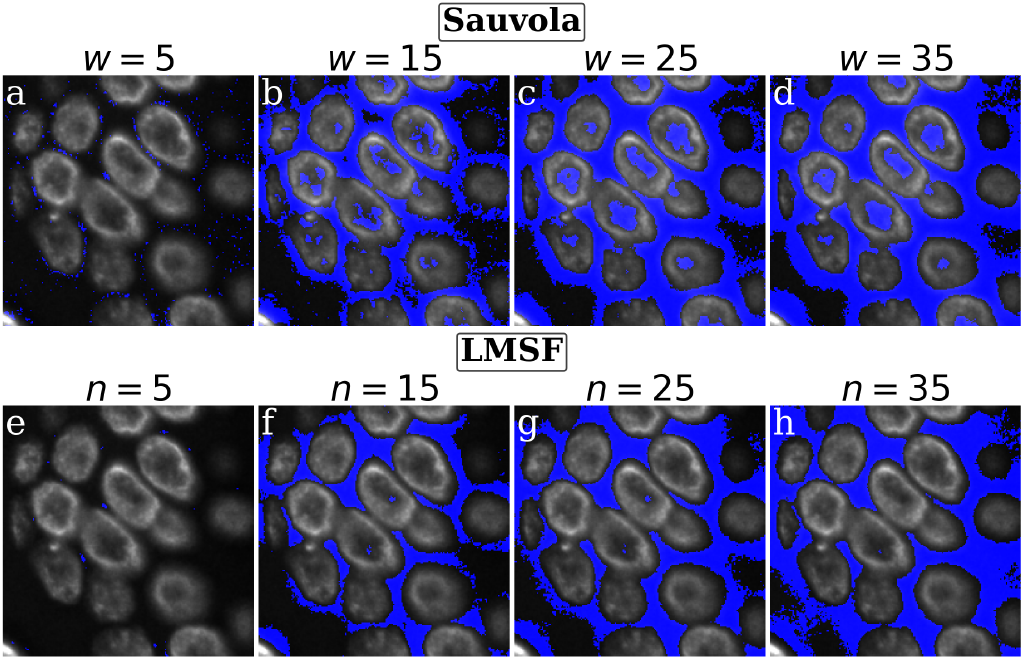
Visual comparison of Sauvola and LMSF methods on a test image containing bright edges and dim middle parts of foreground objects. (a)-(d) Background identified in the test image by Sauvola method corresponding to *k* = 0.2 and *w* = 5, 15, 25, and 35, respectively. (e)(h) LMSF responses (3) to the test image corresponding to *θ* = 0.5 and *n* = 5, 15, 25, and 35, respectively. In both cases, the test image is 200 × 200 fragment of a fluorescence image, where grayscale intensity indicates expression of nucleus marker (Hoechst). Blue color indicates the pixels identified as the background.

## 4 Use Cases

In practical scenarios, fluorescence images of biological tissue have significant diversity in spatial distribution of the foreground markers relative to the image background. Below we present three such scenarios that demonstrate the efficacy of the LMSF method in real-world samples.

### 4.1 Use Case: Large Background

In many practical cases, images of biological tissues have an extensive (large) background, i.e. the number of background pixels is relatively large compared to the number of foreground pixels. An example of such a fluorescence image, along with an illustration of its background identification using our LMSF method, is shown in Fig. 6. The original 1000 × 1000 image with green outlines around foreground objects (nuclei and their clusters) is shown in Fig. 6a, while its binary segmentation indicating the foreground nuclei and background with white and black colors respectively is shown in Fig. 6b. The segmentation was generated using Algorithm 1. As before, we set the threshold parameter *θ* = 0.5. The values of *n*, however, are now given by *N* = {5, 10, 20, 40, 50, 60, 80, 100, 120, 160, 499}. This is because *n*_max_ is now 499. To identify the background between closely located nuclei (short-range and abrupt variations of intensity), we set the averaging parameter to lower values, *n* ∈ {5, 10, 20, 40}. On the other hand, to reliably identify the nonuniform and noisy structure of the extended image background (long-range and gradual variation of intensity), we increased the values of the averaging parameter *n* up to the upper limit, defined by Eq. (4). Fig. 6c shows a Gamma corrected version of the original image with *γ* = 0.5 to highlight the complexity of this image.

**Figure 6.**
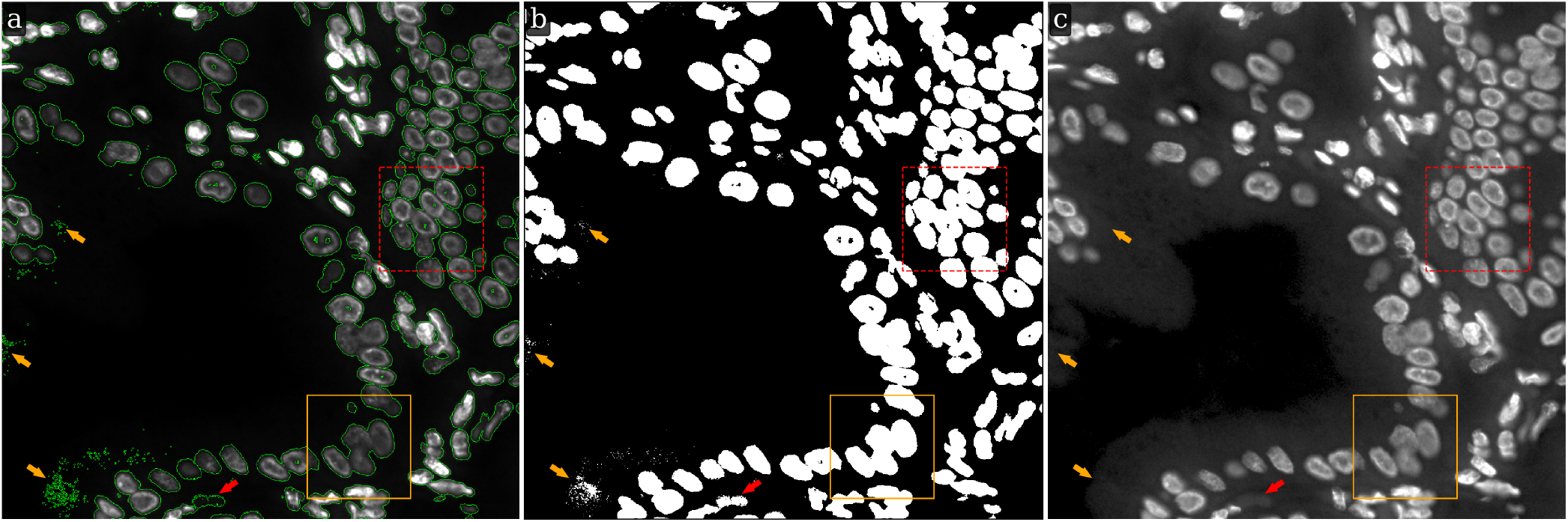
LMSF background identification of an image with a large background. Input image: 1000 × 1000 fluorescence image of a human gallbladder tissue section, where shades of gray indicate expression of nucleus marker (Hoechst). The solid orange (dashed red) bounding box contains the image fragment shown in Figs. 3 and 4 (Fig. 5). (a) The input image with green outlines around foreground objects predicted by the LMSF method with *θ* = 0.5 and *N* = {5, 10, 20, 40, 50, 60, 80, 100, 120, 160, 499}. (b) The binary segmentation of the input image indicating the foreground (background) in white (black) color predicted by the LMSF method. (c) Gamma correction of the input image with *γ* = 0.5 emphasizing the nonuniform and noisy structure of the image background.

As shown in Fig. 6a, despite high image complexity, the LMSF method identified the background almost everywhere, except three small regions (indicated by orange arrows) consisting of point-like objects. All these unidentified background objects have a very small area and are located near the image edges or within background regions with substantial intensity variations as indicated by the orange arrows in Fig. 6c. In addition, these spuriously identified foreground objects are located relatively far from true foreground boundaries, and therefore, can easily be removed using standard image processing methods such as Gaussian blurring combined with Otsu thresholding. Importantly though, the LMSF method accurately identified exact boundaries of all foreground objects including their inhomogeneous and low contrast regions (Fig. 6a). Moreover, having compared the foreground contours and binary mask shown in Fig. 4j with the corresponding contours and mask respectively presented within the orange rectangle in Figs. 6a and b, we infer an important property of the LMSF method, that the wide range of *n* values allows us to detect large background without compromising the boundaries of small foreground objects. This is one more illustration of the LMSF saturation effect mentioned in Section 2.

Moreover, we should note the foreground object indicated by the red arrow in Fig. 6. Looking at Fig. 6a, we see the green contour surrounding some background region and conclude that this object was misidentified as a foreground object by LMSF. However, looking at the Gamma correction of the original image (Fig. 6c) we can see the detailed structure of the image background and identify a solid convex object that has very low contrast. This example demonstrates the high sensitivity and effectiveness of LMSF method.

### 4.2 Use Case: Large Foreground

A different practical case arises in images with densely populated cells and nuclei, i.e. images with an extensive (large) foreground. One such image, and its LMSF method based binarization into background and foreground is shown in Fig. 7. The original 1000 × 1000 image with green outlines around foreground objects (nucleus clusters) is shown in Fig. 7a, while the binary segmentation indicating the foreground objects and background with white and black colors respectively is shown in Fig. 7b. As can be seen, our method is able to correctly identify narrow background regions between large nuclei clusters. As this image does not have large background regions it is natural to limit *n* to a narrow range to avoid compromising foreground shape. Here, we set *θ* = 0.5 and *N* = {5, 10, 20, 40}. Fig. 7c shows the Gamma corrected version of the original image with *γ* = 0.5. It highlights that LMSF method is capable of handling considerable variations in intensity within the dense foreground to precisely identify all narrow background regions and exact boundaries for all foreground nuclei including their low contrast regions.

**Figure 7.**
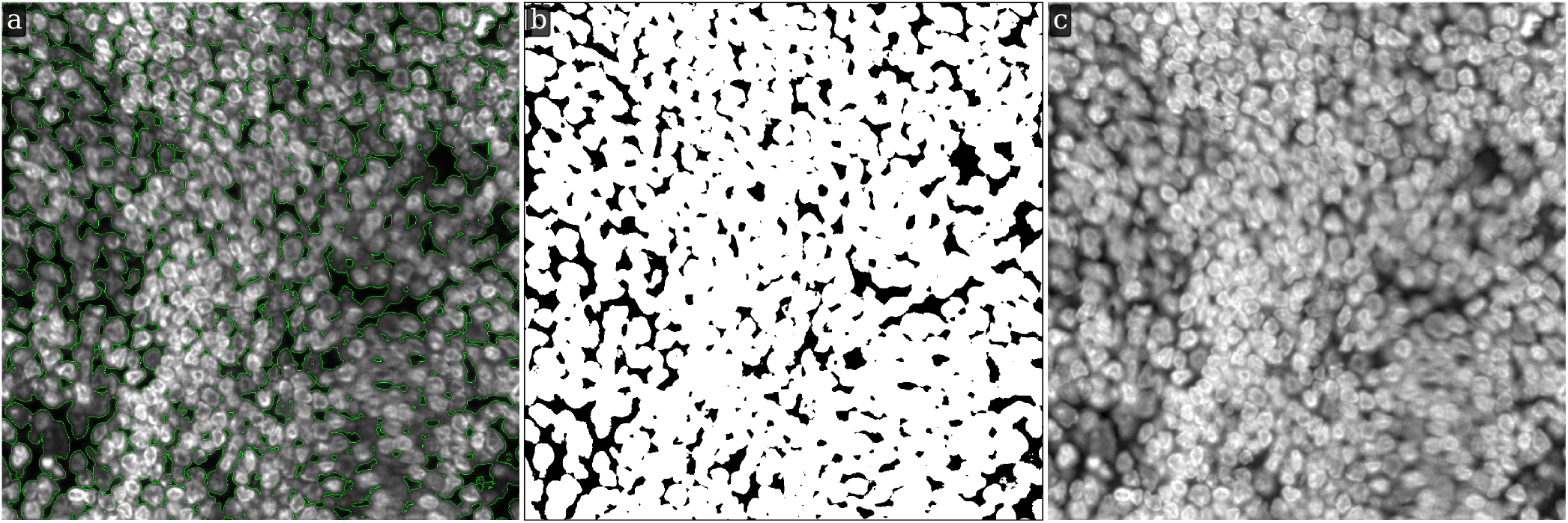
LMSF background identification of an image with a large foreground. Input image: 1000 × 1000 fluorescence image of a human appendix tissue section, where shades of gray indicate expression of nucleus marker (Hoechst). (a) The input image with green outlines around foreground objects predicted by the LMSF method with *θ* = 0.5 and *N* = {5, 10, 20, 40}. (b) The binary segmentation of the input image indicating the foreground (background) in white (black) color predicted by the LMSF method. (c) Gamma correction of the input image with *γ* = 0.5 emphasizing the complex structure of overlapping nuclei (foreground).

### 4.3 Use Case: Fuzzy Foreground

An important practical case arises in images that contain background with intensity values comparable to image foreground. It results in visually unclear (fuzzy) boundaries between background and foreground regions of the images. An example of such image is shown in Fig. 8a. This image shows the expression of Lamin A*/*C, primarily localized to the nucleus membrane. However, due to a variety of experimental reasons, the image capturing Lamin A*/*C expression has a complex foreground and background that are relatively similar and hard to distinguish.

**Figure 8.**
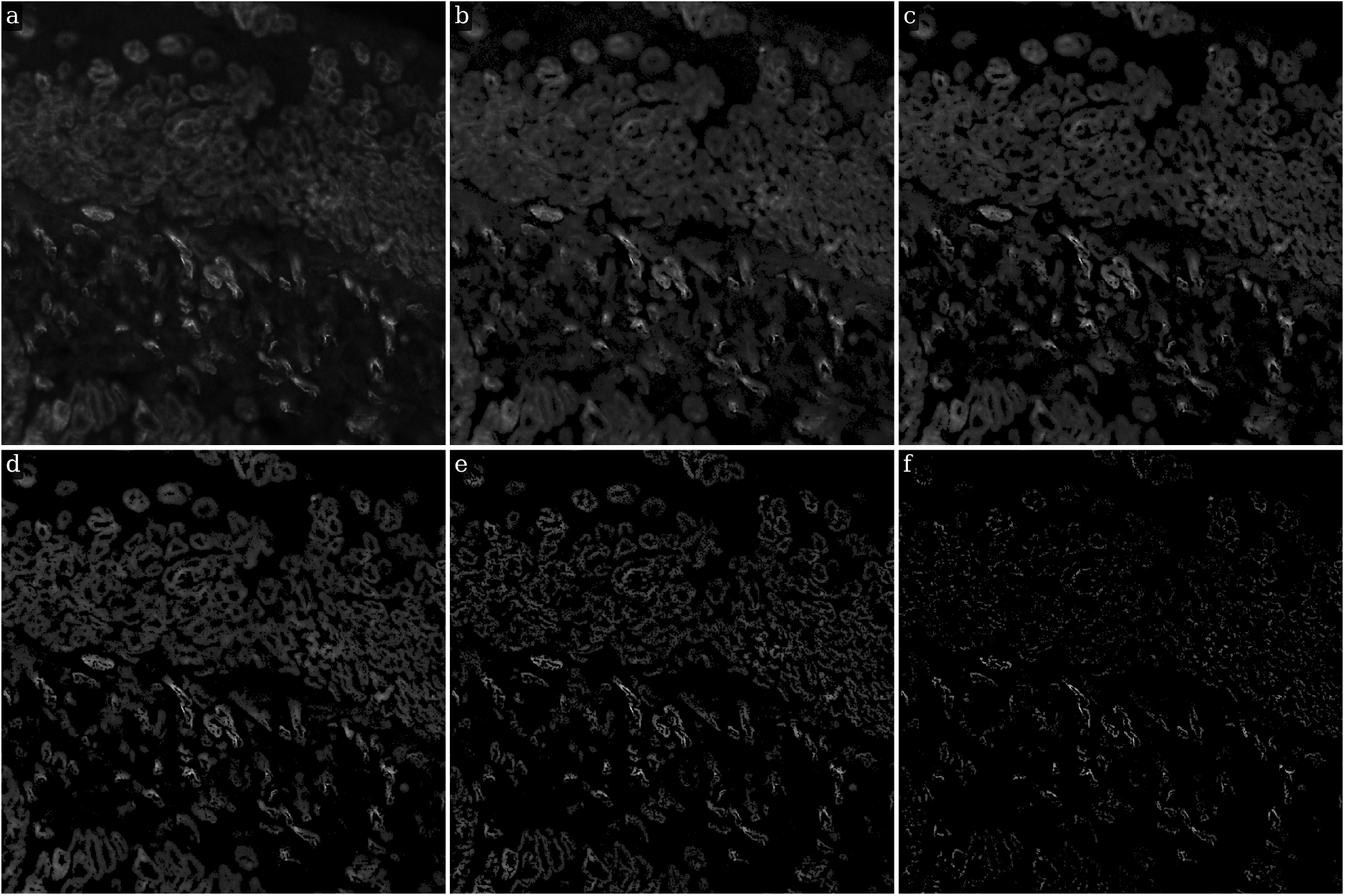
LMSF background identification of an image with a fuzzy foreground. (a) Input image: 1000 × 1000 fluorescence image of tissue section of a human small intestine with adenocarcinoma, where shades of gray indicate expression of nucleus membrane marker (Lamin A*/*C). (b)-(f) The cumulative LMSF responses to the input image calculated using *N* = {5, 10, 20, 40, 80, 160, 499} and setting the threshold parameter *θ* to the following values 0.7, 0.8, 0.9, 1, and 1.1, respectively.

To correctly identify the image background, it is necessary to analyze the spatial variation of image intensity. This analysis can be carried out in the framework of our LMSF method by varying the threshold parameter *θ* and using a fixed range for the averaging parameter *n*. To better capture the contrast between background and foreground we set *N* = {5, 10, 20, 40, 80, 160, 499} focusing more on the detection of image heterogeneity at very short and very long range. Figs. 8b-f show the cumulative LMSF responses to the image shown in Fig. 8a with *θ* = 0.7, 0.8, 0.9, 1, 1.1, respectively. We note that the cumulative LMSF responses calculated for *θ* = 0.9 (Fig. 8d) and *θ* = 1 (Fig. 8e) are closer to effective capturing of the nucleus membrane foreground than the original image (Fig. 8a). This example demonstrates the applicability of LMSF method for background identification in images with fuzzy foreground. It also emphasizes the complexity and nuance of handling fluorescence images of biological tissue sections.

## 5 Applications of LMSF

### 5.1 LMSF based co-localization of molecules profiled using multiplexed imaging

The LMSF method is capable of performing binary segmentation of diverse fluorescence images of different types of molecules. As a consequence, LMSF method can be applied to their multiplexed fluorescence images to characterize their spatial co-localization. For illustration of this idea, we process a 3-channel fluorescence image of adenocarcinoma in a human small intestine (Fig. 9). The first, second, and third channels respectively show the expression of Hoechst (Fig. 9a), Lamin A*/*C (Fig. 9b and Fig. 8a), and Na^+^K^+^ATPase (Fig. 9c). These markers identify the nuclei, nucleus membranes, and cell membranes, respectively. Each channel of the image was independently processed by the LMSF method with the same range for the averaging parameter, *N* = {5, 10, 20, 40, 80, 160, 499} and *θ* respectively set to 0.5, 0.95, and 0.6 for the first, second, and third channels. The resulting binary segmentation masks indicate the spatial localization of Hoechst (Fig. 9d), Lamin A*/*C (Fig. 9e), and Na^+^K^+^ATPase (Fig. 9f). Furthermore, combining these masks into a single image results in 2^3^ classes (Fig. 9h) explicitly indicating the co-localization of the three markers. Co-localization is essential for studying biological mechanisms and this example demonstrates the ability of the LMSF method to aid in such studies. We note that, the LMSF method assumes that image registration of the different multiplexed images has already been performed.

**Figure 9.**
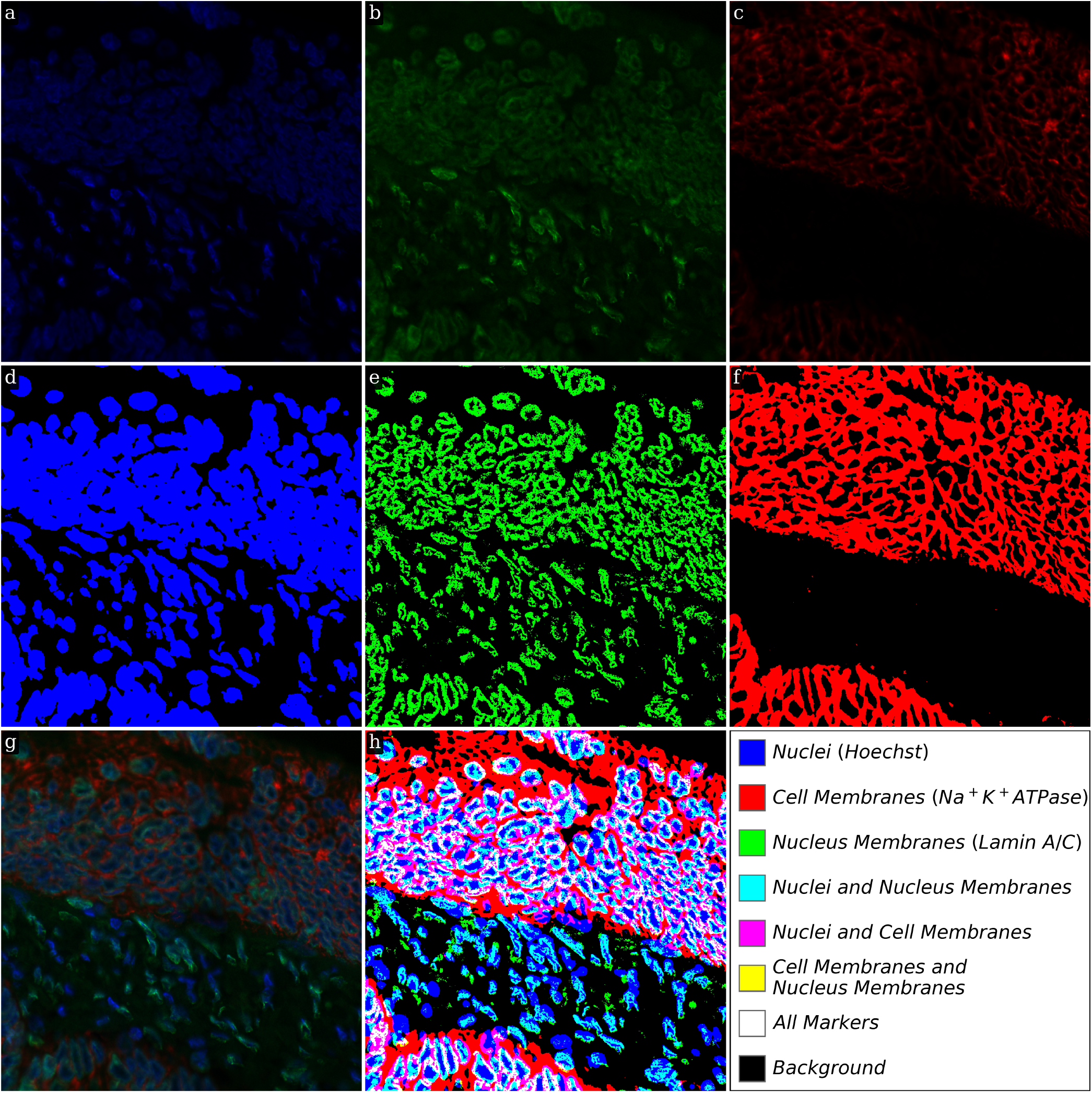
LMSF clustering of 3-channel fluorescence image. Input image: 3-channel image of tissue section of a human small intestine with adenocarcinoma. The sizes of each channel are 1000 × 1000 pixels. (a) First channel: shades of blue indicate expression of nucleus marker (Hoechst). (b) Second channel: shades of green indicate expression of nucleus membrane marker (Lamin A*/*C). (c) Third channel: shades of red indicate expression of cell membrane marker (Na^+^K^+^ATPase). (d) Binary segmentation for nuclei corresponding to the cumulative LMSF response to the first channel calculated with *θ* = 0.5. (e) Binary segmentation for nucleus membranes corresponding to the cumulative LMSF response to the second channel calculated with *θ* = 0.95. (f) Binary segmentation for cell membranes corresponding to the cumulative LMSF response to the third channel calculated with *θ* = 0.6. In all three calculations we put *N* = {5, 10, 20, 40, 80, 160, 499}. (g) Color (RGB) representation for three channels presented in (a)-(c). (h) Logical (Boolean) representation for three binary masks presented in (d)-(f).

### 5.2 Using LMSF method as a denoising step to improve instance image segmentation

Efficient identification of fluorescence image background by LMSF method can help denoise these images and thereby improve accuracy of downstream processes. One such example is improving accuracy of DL methods for image segmentation. We test this proposition using the image shown in Fig. 7, which includes dense nuclei clusters. Fig. 10a shows Mesmer based instance segmentation of individual nuclei, while Fig. 10b shows Mesmer based segmentation preceded by application of LMSF method to accurately identify the background and remove its spurious association with the background. As can be seen, this effective denoising of the image allows Mesmer to generate more accurate and regular nucleus boundaries.

**Figure 10.**
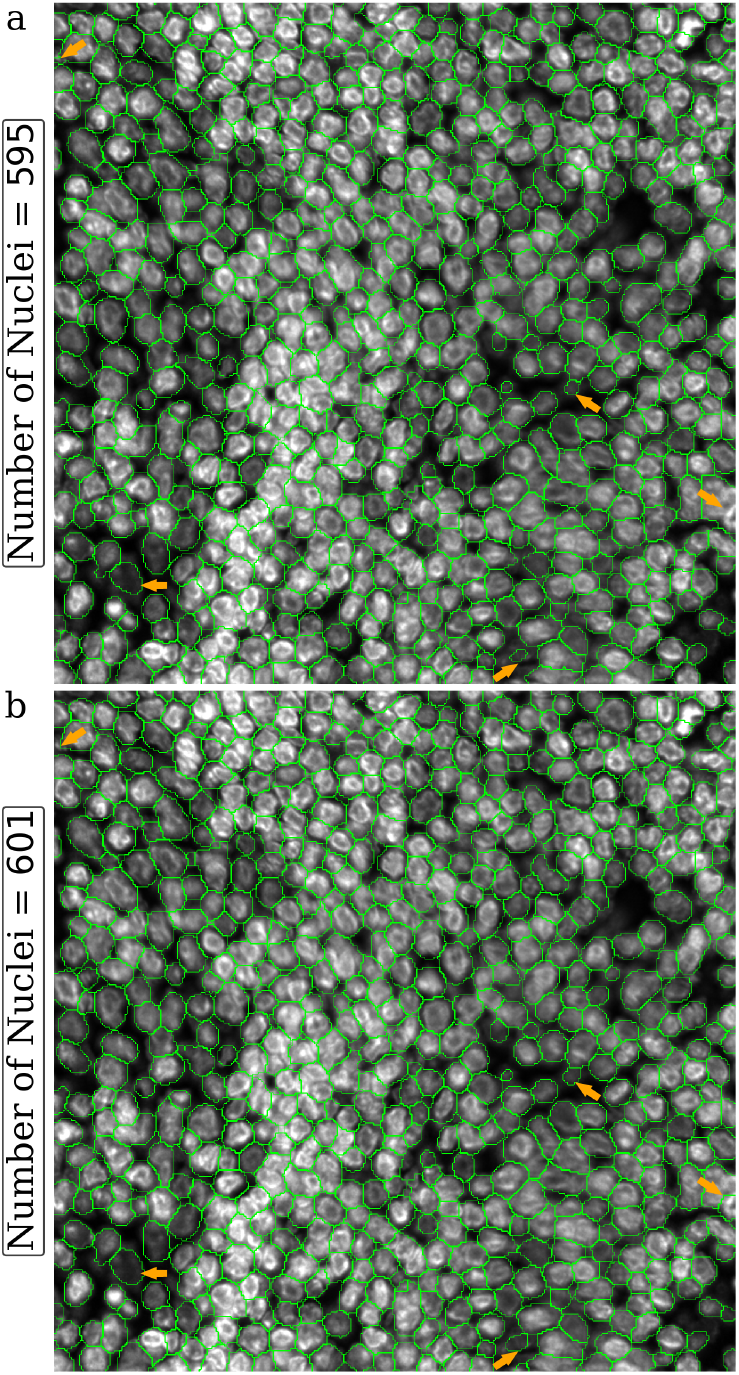
Visual comparison of the effect of LMSF denoising on Mesmer performance of nucleus segmentation. The image is 1000 × 1000 fluorescence image of a human appendix tissue section, where shades of gray indicate expression of nucleus marker (Hoechst). The green lines outline the nuclei predicted by Mesmer with `image_mpp` = 0.16 and `compartment` = *‘whole-cell’* for the unprocessed (a) and LMSF denoised (b) image. To denoise the image we used the cumulative LMSF with *θ* = 0.5 and *N* = {5, 10, 20, 40}. The five orange arrows indicate areas where segmentation has been improved most significantly.

We have also used LMSF method for improving performance of UNSEG, an unsupervised method for segmenting cells and their nuclei in fluorescence images of tissue sections [21].

## 6 Discussion and Conclusions

We have introduced a new local mean suppression filter and its associated method (Algorithm 1) for efficiently and accurately identifying fluorescence image foreground with diverse topologies, low contrast, and inhomogenous signal intensity variations. The LMSF method is able to achieve this by cumulative integration of LMSF responses over multiple scales, using only two LMSF parameters – threshold *θ* and averaging window length *n*.

As a rule of thumb, we observed that *θ* = 0.5 gives excellent results in most cases, except where the boundary between background and foreground is particularly fuzzy, as in Fig. 8a. In such cases, we demonstrated that *θ* can be easily adjusted to perform effective background identification. For images with large foreground and narrow background, the rule of thumb for selecting the range of *n* is as follows: First, we identify the maximal width of the narrow background. This can be an approximation and does not need to be accurate. We set max(*N*) to be half this maximal width. We then continue to recursively divide max(*N*) by 2 and add the result of division to *N* until we reach a reasonable lower limit. For example, in Fig. 7 the maximal background width is approximately 80 pixels. Therefore, we set max(*N*) = 40. Additionally, we added 20, 10, and 5 to *N*. For images with large background, we recommend max(*N*) be computed via (4). The remaining values in *N* should be chosen such that smaller *n* values are emphasized as in Fig. 6.

The testing of LMSF method on diverse and complex fluorescence images and its comparison with robust unsupervised and state-of-the-art DL methods confirms its high performance quality and effectiveness in background identification over these methods. Importantly, LMSF method can be applied to accurately capture the distribution of any molecule of interest and is not limited to those associated with cell membranes and nuclei. Additionally, although not discussed here, LMSF method can also be used to process non-biological grayscale images. In conclusion, we have introduced a simple filter and an easy to use method for identifying background of fluorescence images that demonstrates high accuracy and low implementation cost compared to the current state of the art. We have also included diverse use cases of its applications in real-world scenarios.

## Code availability

The Python implementation of LMSF for processing of signal sequences [Eq. (2)] and grayscale images [Eq. (3) and Algorithm 1] is available at https://github.com/uttamLab/LMSF.

## Data availability

The fluorescence images used in this paper have been taken from the GIT dataset [21], which is available at https://doi.org/10.7303/syn61804540.

## Acknowledgments

This work is supported by the National Cancer Institute (NCI) at the National Institutes of Health (NIH), Grant Number 1R21CA289340-01A1.

